# Live cell biosensors based on the fluorescence lifetime of environment-sensing dyes

**DOI:** 10.1101/2022.02.08.479035

**Authors:** Brian P. Mehl, Li Li, Elizabeth Hinde, Bei Liu, Christopher J. MacNevin, Chia-Wen Hsu, Enrico Gratton, Klaus M. Hahn

**Affiliations:** Department of Pharmacology, University of North Carolina at Chapel Hill, Chapel Hill, NC 27599; Laboratory for Fluorescence Dynamics, Department of Biomedical Engineering, University of California at Irvine, Irvine, CA 92617

**Keywords:** biosensor, Cdc42, FLIM, solvatochromic, dye, merocyanine, phasor, fluorescence

## Abstract

Most biosensors used with fluorescence lifetime imaging (FLIM) have been based on fluorescence resonance energy transfer (FRET). Here we examined the capabilities of FLIM biosensors based on environment-sensing dyes. We screened merocyanine dyes to find an optimal combination of environment-induced FLIM changes, photostability and brightness at wavelengths suitable for live cell imaging. A biosensor reporting conformational changes of endogenous Cdc42 protein was tested in vitro and in live cells. In addition to the known advantages of FLIM (quantitation independent of photobleaching, intracellular biosensor distribution, and excitation intensity), the spectral properties of the merocyanine-FLIM biosensor (mcFLIM) provided enhanced sensitivity at low activation levels (<10%) where FRET is typically least sensitive. We leveraged these properties and the phasor representation of FLIM to determine the specific concentration of activated Cdc42 across the cell.

## INTRODUCTION

Fluorescent biosensors have been a powerful tool for untangling signaling behavior in living cells (Greenwald et al., 2018; Komatsu et al., 2011; Machacek et al., 2009; Marston et al., 2020; Pertz et al., 2006; Yasuda et al., 2006). Most often they have been based on fluorescence resonance energy transfer (FRET), with readouts quantified using ratiometric fluorescence intensity measurements. Fluorescence lifetime imaging microscopy (FLIM) can provide important advantages over intensiometric measurements, for FRET and for other biosensor approaches (Bastiaens and Squire, 1999; Hinde et al., 2012, 2013). FLIM readouts are independent of fluorophore concentration, excitation intensity, and photobleaching. This is particularly relevant when imaging within living systems, where excitation is affected by tissue refraction, or where variations in biosensor concentration affect intensity. With intensity readouts, artefactual changes due to incomplete protein folding, variable expression or photobleaching cannot be readily separated from the salient FRET changes, because all occur at the same wavelengths. FLIM not only distinguishes FRET from artefacts, but does so using a single fluorophore, simplifying multiplexed imaging. (Berney and Danuser, 2003; Datta et al., 2020; Elangovan et al., 2003; Malacrida et al., 2021).

For biosensors, FLIM has been used almost entirely to image FRET (Berezin and Achilefu, 2010; Hinde et al., 2012, 2013; Ng et al., 1999; Orthaus et al., 2009; Wallrabe and Periasamy, 2005) but biosensors based on environment-sensing dyes have distinct advantages over FRET (Yasuda, 2006; Yasuda et al., 2006). Such biosensors have recently become more accessible through attachment of dyes to proteins within living cells, eliminating the need for electroporation or other cumbersome techniques to load cells. Environment-sensing dyes have either been attached to target proteins where they are affected by conformational changes (Garrett et al., 2008; Hahn et al., 1992; Nomanbhoy et al., 1996), or have been attached to “affinity reagents” that bind selectively to the activated conformation of the target, causing a spectral change in the attached dye(Gulyani et al., 2011; Nalbant et al., 2004). Affinity reagents have been based on proteins, peptides, or even small molecules(Gulyani et al., 2011; Konze et al., 2013; Nalbant et al., 2004) (**Figure 1a**). They can report the activation of endogenous, unmodified targets. Dye-based biosensors can produce much brighter emission than FRET because the dyes are directly excited (Gulyani et al., 2011; MacNevin et al., 2013; Nalbant et al., 2004). To date, environment-sensing dyes have been monitored in cells primarily through ratiometric fluorescence intensity changes (Gulyani et al., 2011; MacNevin et al., 2013, 2016, 2019; Nalbant et al., 2004); Biosensors based on the FLIM of such dyes remain largely unexplored (Berezin et al., 2009; Klymchenko, 2017).

**Figure 1.**
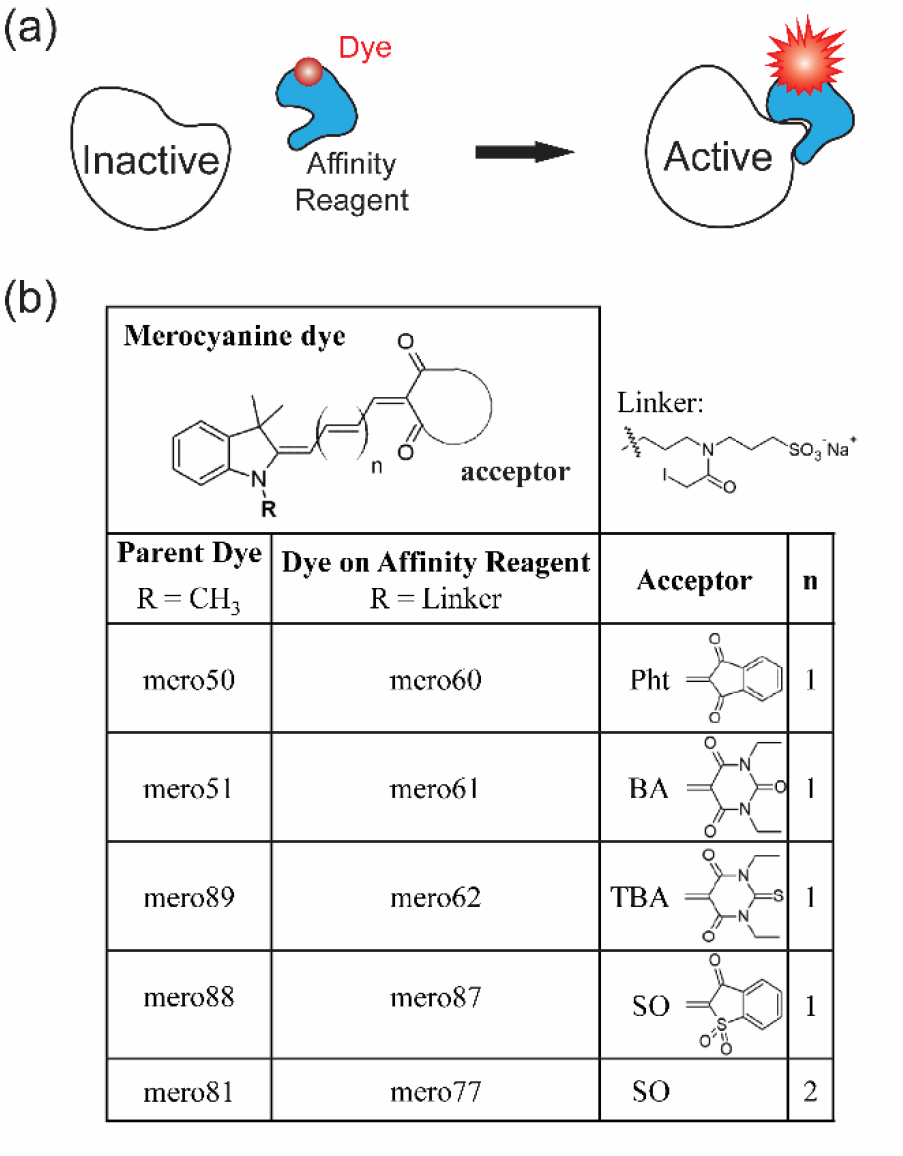
(**a**) The dye-based Cdc42 biosensor, consisting of an affinity reagent that specifically binds to the active conformation of Cdc42 and a covalently-attached environment-sensitive dye (red). Upon Cdc42 activation, the affinity reagent binds to Cdc42, altering the dye’s local environment and affecting optical properties including brightness and fluorescence lifetime. (**b**) Parent dyes modified with a cysteine-reactive linker (upper right) and used to construct biosensors. Abbreviations: phthalimide (Pht), barbituric acid (BA), thiobarbituric acid (TBA), and benzothiophene (SO). **n** indicates number of double bonds, as shown in the structure at top.

Here we make a FLIM-based biosensor by attaching a merocyanine dye with environment-sensitive lifetime to an affinity reagent that selectively binds the activated conformation of Cdc42. Dyes were screened for optimal fluorescence and solvent-sensitive lifetime, and the dye with the best combination of brightness and lifetime response to Cdc42 binding was used in the final biosensor. The affinity reagent was derived from previous biosensors that used dyes undergoing ratiometric intensity changes (MacNevin et al., 2013, 2016, 2019; Nalbant et al., 2004) **(Figure 1a)**. The final FLIM biosensor provided sensitive readouts of Cdc42 activity in living cells. Importantly, the merocyanine dyes became brighter as the biosensor bound Cdc42, so the bound biosensor contributed more to the overall lifetime. This enhanced detection of low Cdc42 concentrations, where FRET is least sensitive. This should be especially valuable for GTPases, which can initiate cell behaviors when less than 10% of the protein population is active (Boulter et al., 2010; Jennings and Knaus, 2014). By adapting an approach used for calcium biosensors we were able to determine the actual concentration of activated species, not just the relative distribution of activity. Our results identify dyes for FLIM biosensors, show how they can be used to report activity of an endogenous target protein, demonstrate the advantages of FLIM, and show how to quantify the concentration of active species.

## MATERIALS AND METHODS

### Dye synthesis and protein conjugation

Dyes were synthesized as previously described (MacNevin et al., 2013), except for the new dyes **81** and **77**, whose synthesis is described below. Production of the WASP domain used as the affinity reagent, and its conjugation with dyes, was as described in Macnevin et al. (MacNevin et al., 2013), a modification of the procedure in Nalbant et al. (Nalbant et al., 2004).

2,3,3-Trimethylindolenine was purchased from TCI America. *N*-(5-Anilino-2,4-pentadienylidene)aniline hydrochloride was purchased from Acros Organics. All other reagents were purchased from Sigma-Aldrich. The following compounds were prepared according to published procedures: benzo[b]thiophen-3(2H)-one 1,1-dioxide(Regitz and Stadler, 1965), 1,2,3,3-tetramethylindolium iodide (Wang et al., 2003), and 2,3,3-trimethyl-1-(3-((3-sulfopropyl)amino)propyl)indolinium bromide (MacNevin et al., 2013). Reactions were run using anhydrous solvents. All operations with dyes were performed under dim light. UV-Visible spectra were recorded on a Hewlett-Packard 8453 diode array spectrophotometer. Fluorescence spectra were obtained on a Spex Fluorolog 2 spectrofluorometer. Reverse phase high performance liquid chromatography was performed on a Shimadzu Prominence system with a 250 x 21.2 mm, 15 micron Phenomenex C18 preparative column and elution at 8 mL/min with a gradient of 10% solvent B (H_2_O/ACN 5:95, TFA 0.05%) 90% solvent A (H_2_O/ACN 95:5, TFA 0.05%) for 2 min, increasing to 90% solvent B over 30 min and held for a total of 45 min. NMR spectra were taken on a Varian Mercury-300 or Inova-400 spectrometer using deuterated solvents purchased from Cambridge Isotope Laboratories. Reference peaks of 7.26 ppm (CDCl_3_), 2.50 ppm (DMSO-d_6_), and 2.92 ppm (DMF-d_7_) were used for ^1^H spectra. Low resolution mass spectra were collected on an Agilent MSD Ion Trap mass spectrometer with direct infusion. High resolution mass spectra were obtained on an Agilent 6520 Accurate-Mass Q-TOF LC-MS.

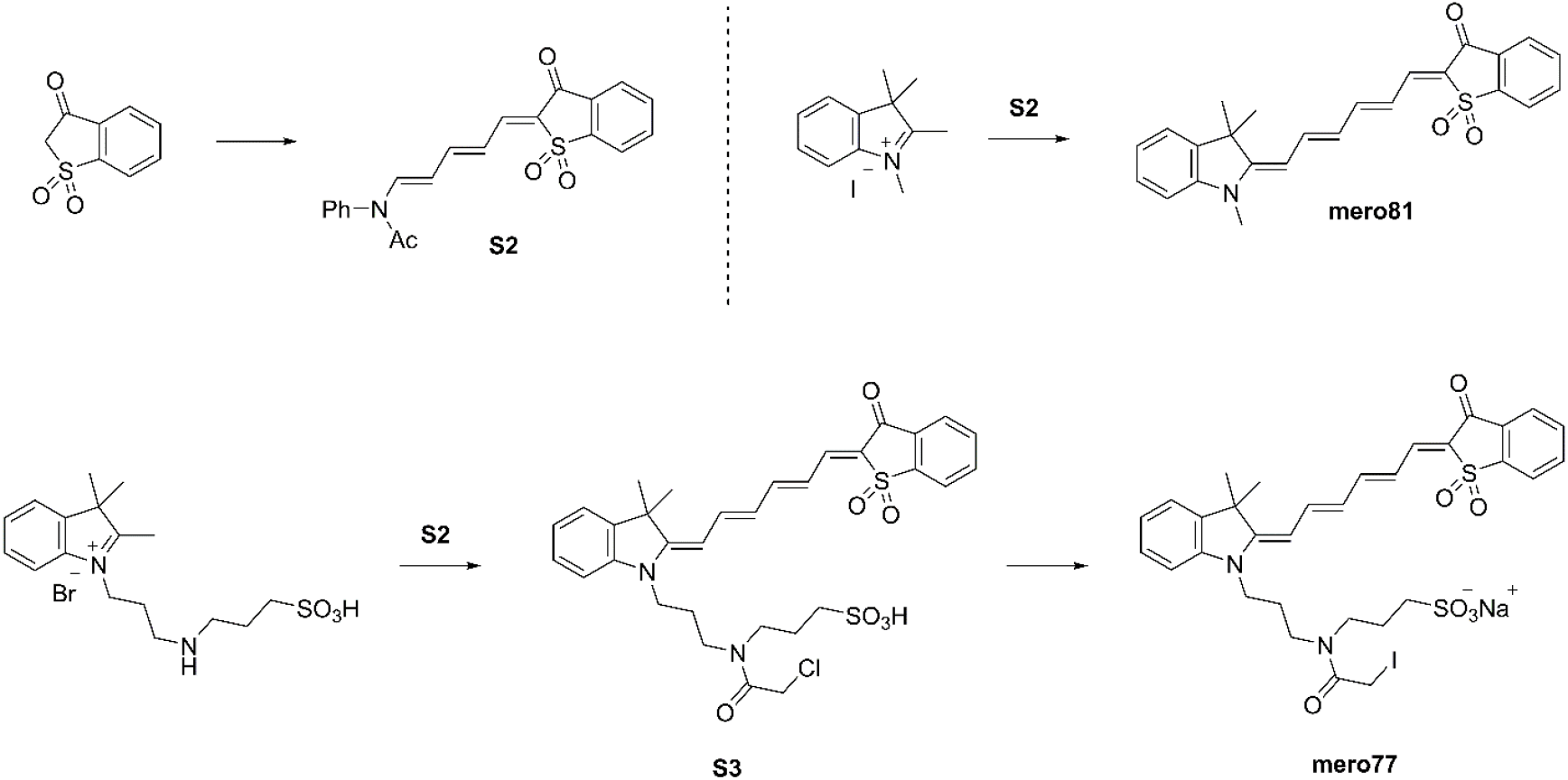

#### *N*-((1*E*,3*E*,5*E*)-5-(1,1-dioxido-3-oxobenzo(*b*)thiophen-2(3*H*)-ylidene)penta-1,3-dien-1-yl)-*N*-phenylacetamide (S2)

A mixture of the benzo[b]thiophen-3(2H)-one 1,1-dioxide (2.73 g, 15.0 mmol) and N-(5-anilino-2,4-pentadienylidene)aniline hydrochloride (5.98 g, 21.0 mmol) were diluted in acetic anhydride (25 mL). The mixture was heated to reflux under argon for 2 h and cooled to room temperature. The resulting precipitate was filtered, washed with ethyl acetate, and dried under vacuum. The product was re-crystalized from isopropanol to give 4.16 g dark brown solid (73% yield). ^1^H NMR (400 MHz, CDCl_3_) δ 8.14 (d, 1H, *J* = 13.8 Hz), 8.01 (d, 1H, *J* = 7.6 Hz), 7.92 (d, 1H, *J* = 7.7 Hz), 7.82 (t, 1H, *J* = 7.4 Hz), 7.75 (t, 1H, *J* = 7.6 Hz), 7.53-7.64 (m, 4H), 7.19 (d, 2H, *J* = 7.7 Hz), 7.12 (d, 1H, *J* = 11.2 Hz), 6.79 (t, 1H, *J* = 13.6 Hz), 5.34 (t, 1H, *J* = 13.6 Hz), 1.96 (s, 3H). MS-ESI *m/z* 402.0 ([M + Na]^+^ requires 402.1).

#### Mero81

1,2,3,3-Tetramethylindolium iodide (0.15 g, 0.50 mmol), compound **S2** (0.21 g, 0.55 mmol), and sodium acetate (0.053 g, 0.65 mmol) were dissolved in methanol (2.0 mL) and heated to reflux for 10 min. The reaction mixture was cooled to room temperature, concentrated via rotary evaporation, and purified by silica gel chromatography using dichloromethane and ethyl acetate. The product was isolated as 0.136 g blue-olive solid (65% yield). ^1^H NMR (400 MHz, CDCl_3_) δ 8.00 (d, 1H, *J* = 7.6 Hz), 7.94 (d, 1H, *J* = 7.5 Hz), 7.66-7.81 (m, 3H), 7.38-7.47 (m, 1H), 7.21-7.28 (m, 3H), 7.03 (t, 1H, *J* = 7.4 Hz), 6.93 (t, 1H, *J* = 13.32 Hz), 6.82 (d, 1H, *J* = 6.8 Hz), 6.38 (t, 1H, *J* = 12.7 Hz), 5.64 (d, 1H, *J* = 12.8 Hz), 3.31 (s, 3H), 1.63 (s, 6H). MS-ESI *m/z* 418.3 ([M + H]^+^ requires 418.2).

#### 3-(2-chloro-*N*-(3-((*E*)—((2*E*,4*E*,6*E*)-6-(1,1-dioxido-3-oxobenzo(*b*)thiophen-2(3*H*)-ylidene)hexa-2,4-dien-1-ylidene)-3,3-dimethylindolin-1-yl)propyl)acetamido)propane-1-sulfonic acid (S3)

2,3,3-Trimethyl-1-(3-((3-sulfopropyl)amino)propyl)indolinium bromide (0.225 g, 0.50 mmol), chloroacetic anhydride (0.427 g, 2.50 mmol), and sodium acetate (0.123 g, 1.50 mmol) were diluted in DMF (5.0 mL). The solution was stirred at room temperature for 15 min, followed by addition of compound **S2** (42.3 mg, 0.11 mmol). The reaction mixture was stirred at room temperature under argon for 2 h. The majority of DMF was removed via azeotropic distillation with toluene under reduced pressure. The residue was purified by gradient elution on a silica gel column using a 0 to 10% methanol in dichloromethane gradient. The product was isolated as 0.049 g dark blue solid (15% yield). ^1^H NMR (400 MHz, DMSO-d_6_) δ 8.01 (d, 1H, *J* = 7.2 Hz), 7.49 (d, 1H, *J* = 7.4 Hz), 7.26-7.36 (m, 1H), 7.21 (d, 1H, *J* = 7.8 Hz), 7.11 (t, 1H, *J* = 7.3 Hz), 6.46-6.58 (m, 1H), 6.04-6.20 (m, 1H), 4.39 (s, 2H), 4.26-4.32 (m, 2H), 3.90-4.00 (s, 2H), 2.30-2.50 (m, 4H), 1.80-1.95 (m, 2H), 1.61 (s, 6H). HR-MS (Q-TOF) *m/z* 657.1490 ([M - H]^-^ requires 657.1501).

#### Mero77

Compound **S3** (22.7 mg, 0.036 mmol) and sodium iodide (53.9 mg, 0.36 mmol) were diluted in 1:1 chloroform/methanol (1.0 ml) and refluxed for 3 h under nitrogen. The reaction mixture was concentrated, diluted in a minimum amount of methanol, and submitted to reverse phase HPLC purification. Product fractions were combined and lyophilized to give a dark blue solid. ^1^H NMR (DMF-d_7_), 7.87-8.08 (m, 4H), 7.80 (t, 1H, *J* = 12.9 Hz), 7.62-7.64 (m, 1H), 7.58-7.60 (m, 1H), 7.53 (d, 1H, *J* = 7.8 Hz), 7.32-7.45 (m, 2H), 7.16 (t, 1H, *J* = 7.2 Hz), 6.96 (t, 1H, *J* = 13.5 Hz), 6.53-6.65 (m, 1H), 6.29 (d, 1H, *J* = 13.5 Hz), 4.07-4.25 (m, 4H), 3.45-3.3.70 (m, 4H), 2.71 (t, 2H, *J* = 6.6 Hz), 2.01-2.16 (m, 4H), 1.69-1.71 (m, 8H). MS-ESI *m/z* 751.1 ([M + H]^+^ requires 751.1).

### Lifetime Measurements

*In vitro* and *in vivo* fluorescence lifetimes were collected on a modified laser scanning confocal microscope (Olympus FV1000) equipped with a UPLFL40X, 1.30 NA objective. The merocyanine dyes were excited at 594nm using a pulsed supercontinuum laser source (Fianium) at 20 MHz in combination with an acousto-optic tunable filter (AOTF) (Crystal Technologies). The images were obtained with 256 x 256, 20μs/pixel, 1.660 s/frame acquisition settings. Lifetime images were averaged over 35 frames for a total acquisition period of 58 s. The emission was detected with an external GaAsP photomultiplier detector (Hamamatsu: H722P-40). An emission bandpass filter (Semrock: FF-02-628/40-25) was placed before the PMT to reject unwanted excitation light. A digital frequency domain approach was used to acquire lifetime measurements (ISS: A320 FastFlim box) (Colyer et al., 2008). Calibration of the system was performed by measuring Texas red, which has a known lifetime of 4.2 ns in water. Processing was done with custom scripts developed in Matlab^®^.

The fluorescence decay of each merocyanine dyes was acquired with a time-correlated single photon counting fluorometer based on an ISS Koala automated sample compartment, supercontinuum laser source (Fianium), Hamamatsu photomultiplier tube, and Becker & Hickl SPC-144 board. For each acquisition, the bin time was 24.43ps and points were integrated over 60 seconds. Depending on the spectral properties of each merocyanine dye the Fianium laser was tuned to either 570nm or 600nm while emission was collected using a 600/25 nm or 625/25 nm bandpass filter placed in the front of the detector. The instrument response was calibrated by measuring the instantaneous response from an aqueous dispersion of glycogen.

### Calculation of Active Cdc42 Concentration

The following is based on a similar derivation previously published for the calcium sensor, CG5N (Celli et al., 2010). The following description has been included for completeness. The free, ***P_f_*** and bound, ***p_b_*** forms of the biosensors have unique locations within the phasor plot, defining characteristic lifetimes for both states. Utilizing the property of linear combinations, a phasor ***p***, which contains a portion of free and bound biosensor, will be located somewhere along a line connecting the two states. This phasor position is given by the following equation.

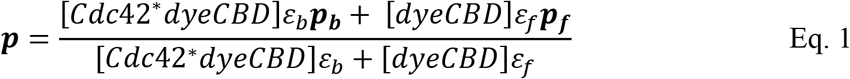

*ε_f_* and *ε_b_* correspond to the relative brightness of the free (*dyeCBD*) and bound (*Cdc42*dyeCBD*) states of the biosensor, respectively. The asterisk designates that Cdc42 is in its active, GTP-bound conformation. With knowledge of these phasor locations along with the relative brightness of the two states, the fraction of bound biosensor, *Fb* can be calculated.

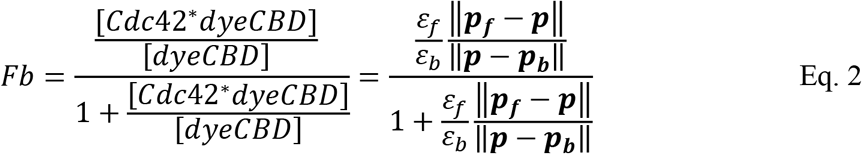

To determine the concentration of active Cdc42, the dissociation constant, *K_d_* must be determined. Mathematically, this is presented through the law of mass action.

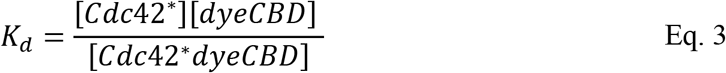

When Eq. 3 is inserted into Eq.2 and after some algebraic reorganization, the concentration of active Cdc42 is given by Eq. 5. It is proportional to the dissociation constant and dependent on the fraction of bound biosensor.

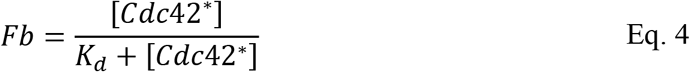

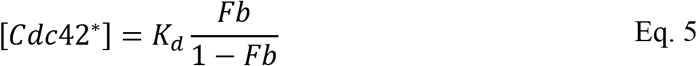

The dissociation constant, and ratio of free to bound brightness were previously published for **mero87-CBD**; 186 ± 40 nM and 1/8.9 respectively (MacNevin et al., 2013). Pseudo-colored images of activated Cdc42 concentration were generated by first determining the fraction of bound biosensor for every pixel. This was achieved by projecting the measured phasor ***p*** for each pixel on the line connecting ***p_f_*** and ***p_b_*** (Eq. 2). After this, the concentration was determined using Eq. 5, with knowledge of the dissociation constant.

## RESULTS & DISCUSSION

### Merocyanine dyes

Merocyanine dyes that are bright and fluoresce in the red portion of the spectrum can respond to changes in their solvent environment(MacNevin et al., 2013, 2016, 2019; Toutchkine et al., 2003). They were therefore tested for FLIM changes that could be used in biosensors. Merocyanine dyes are characterized by electron donor and acceptor moieties linked together through conjugation. Previous studies showed that the local solvation environment could result in large changes in fluorescence intensity as well as shifts in excitation and emission maxima(Han et al., 2003; Liu et al., 2004; Toutchkine et al., 2007). It was observed that an indolenine donor could be combined with different strong electron acceptors to produce merocyanines with a useful compromise between brightness, photostability and environment-sensitive fluorescence(MacNevin et al., 2013, 2016, 2019; Toutchkine et al., 2003, 2007). These acceptors included phthalimide (Pht), barbituric acid (BA), thiobarbituric acid (TBA) and benzothiophene (SO). The dyes used in this study were based on indolenine linked to one of these bases via a conjugated chain **(Figure 1b)**. Parent dyes were assessed for fluorescence properties, and the fluorophores were then derivatized with a cysteine-selective linker for attachment to the affinity reagent **(Figure 1)**.

The dyes were screened for the sensitivity of their fluorescence lifetimes to environment. **Figure 2** shows the measured phase lifetimes, τ_p_ for each of the parent dyes prior to attachment of the cysteine-reactive linker. Lifetime was measured in three solvents with different polarity and hydrogen bonding strengths (Hansen, 2000). The parent merocyanine dyes were **mero50, mero51, mero89** and **mero88** (MacNevin et al., 2013), all of which contain an indolenine donor, but different acceptors (Pht, BA, TBA, and SO). A previously unpublished dye, **mero81** was also evaluated. Its extended polyene system produced red-shifted absorption and emission spectra that could be valuable in multiplexed imaging applications.

**Figure 2.**
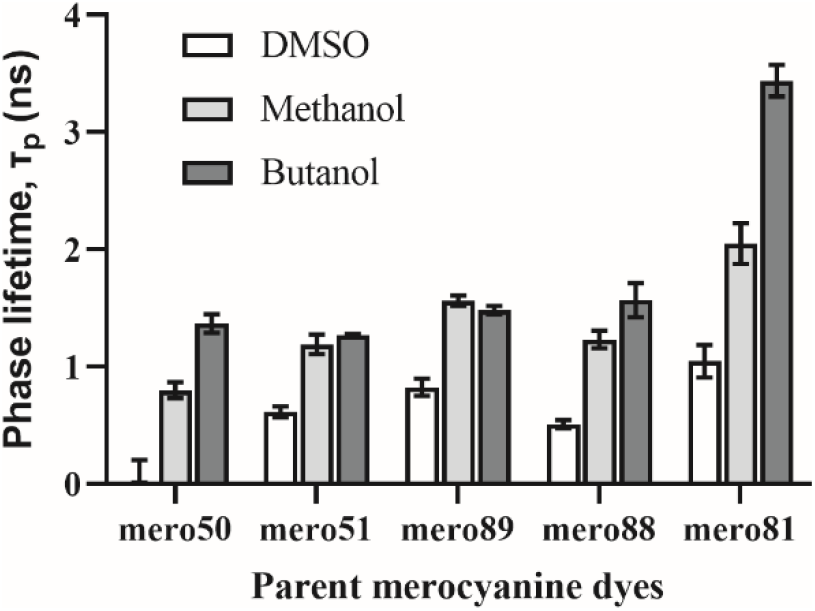
Phase lifetime measurements of parent merocyanine dyes in DMSO, methanol, and butanol (n = 3).

All parent dyes had lifetimes in the nanosecond range, which lend themselves to current lifetime instrumentation (Becker, 2012). The shortest lifetimes were observed when the parent dyes were measured in the strongly polar, relatively weak hydrogen bonding solvent DMSO. In contrast, the longest lifetimes were observed in the least polar hydrogen bonding solvent, butanol. Response to environment can be seen as resulting from differential solvent interaction with specific resonance structures(Blanchard-Desce et al., 1995; Bublitz et al., 1997; Würthner et al., 1997, 2002) or electron distributions(Han et al., 2003; Liu et al., 2004; Toutchkine et al., 2003, 2007).

### *In Vitro* assay of biosensor response

The extent of solvent-dependent changes in FLIM were promising for biosensor construction, but it was hard to predict how Cdc42 binding of the affinity reagent would change the dyes’ environment and affect FLIM. Therefore, each dye was tested in an actual biosensor. A water-soluble, cysteine-reactive side chain was attached to each dye for covalent coupling to the single cysteine of the affinity reagent. This was based on the approach previously used to attach intensiometric dyes to the same cysteine (Nalbant et al., 2004). The affinity reagent was based on the Cdc42 binding domain of Wiskott Aldrich Sydrome Protein (WASP) (MacNevin et al., 2013; Nalbant et al., 2004). It has been shown that this domain binds selectively to the activated, GTP-bound conformation of Cdc42 (Abdul-Manan et al., 1999). In-vitro assays were performed to measure the difference in lifetime between bound and unbound biosensor. The final products were named mcFLIM biosensors, for merocyanine FLIM, with a number denoting the dye (e.g. mcFLIM 87 for the affinity reagent labeled with dye mero87).

The phasor approach is a powerful way to visualize lifetime changes without the aid of complex fitting routines (Colyer et al., 2008; Digman et al., 2008; Malacrida et al., 2021). Displayed in **Figure 3a** is a phasor plot of **mcFLIM 87** in the presence and absence of a constitutively active Cdc42 mutant (Q61L). The free and bound forms of the biosensor had unique phasor locations within the phasor plot, defining characteristic lifetimes for each state.

**Figure 3.**
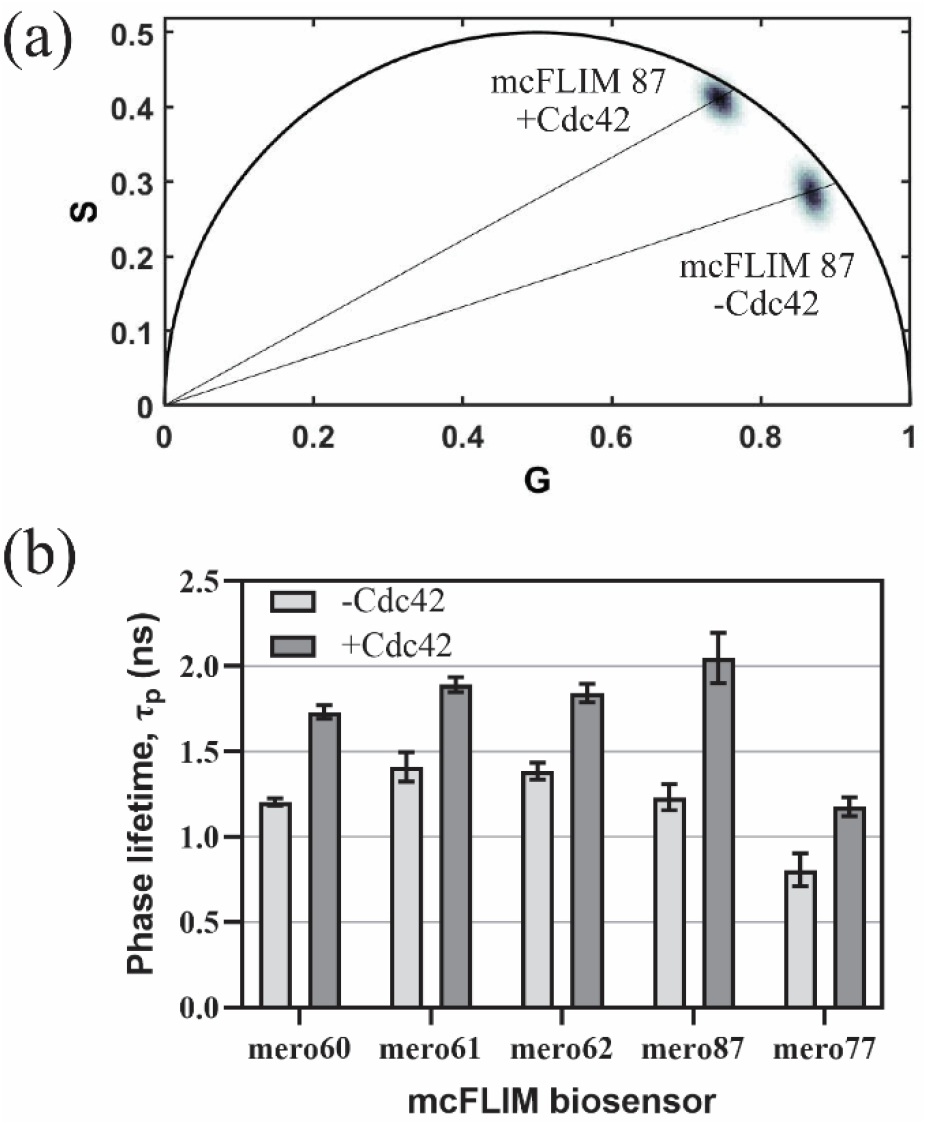
*In vitro* assay for lifetime changes associated with Cdc42 binding. (**a**) Phasor histogram of **mcFLIM 87** biosensor (400 nM) in the presence and absence of 12 μM constitutively active Cdc42 mutant (Q61L). (**b**) Bar graph displaying phase lifetimes of biosensors made with the indicated dyes (n =3).

Utilizing the property of linear combination, a phasor which contains a portion of free and bound biosensor will be located somewhere along a line connecting the two distinct states. We observed that upon biosensor binding to Cdc42, the lifetime of all tested biosensors increased **(Figure 3b)**. Additionally, the phasors did not lie on the “universal circle” (Digman et al., 2008; Hinde et al., 2012; Malacrida et al., 2021), indicating complex decay kinetics. Fortunately, dye-based biosensors, unlike FRET-based biosensors, do not require monoexponential decay kinetics to quantify protein activity (Datta et al., 2020; Orthaus et al., 2009).

All Cdc42 biosensors exhibited an appreciable change in lifetime. **mcFLIM 87** showed the largest difference between the free and bound conformations. Although the dye mero81 performed the best in the pure solvent tests **(Figure 2)**, the biosensor derived from it (**mcFLIM 77**) showed a relatively weak response to Cdc42 binding. We chose to pursue **mcFLIM 87** because of its strong response to Cdc42.

### Detecting low concentrations of activated proteins

All the biosensors increased their emission intensity upon activation, a property of the merocyanine dyes used. The overall lifetime was therefore substantially impacted by even a small fractional contribution from the active form. The FLIM biosensors provided enhanced sensitivity in precisely the concentration range where FRET biosensors are least sensitive. In FRET the directly excited fluorophore (the donor) decreases intensity upon activation. **Figure 4** shows the fractional contribution of the activated form to the overall lifetime, for dye and FRET-based biosensors. A biosensor designed to produce increased FRET when the target is in the active conformation (as most are) has a poor fractional contribution from the quenched state at low protein activity levels (5% fractional contribution at 10% protein activity, **Figure 4** solid red line). Biosensors can be designed so that activation decreases FRET, increasing donor emission, but there are few examples of this used for FRET-FLIM (Lee et al., 2009; Marston et al., 2020; Stirnweiss et al., 2013). A dye-based biosensor with a free to bound brightness ratio of 0.1, like **mcFLIM 87**, has a large, 50% fractional contribution from the bound state at 10% activity level (**Figure 4** blue dotted line). To produce a similar lifetime change, a FRET-based biosensor would need to achieve 90% FRET in the inactive conformation.

**Figure 4.**
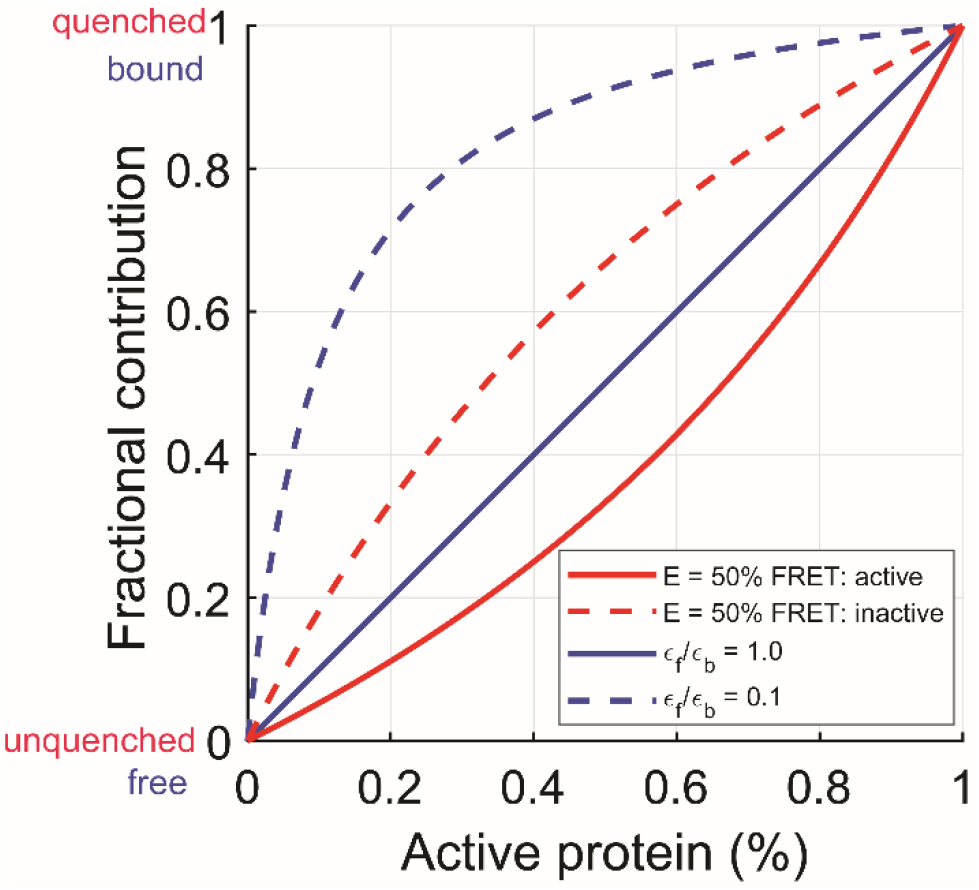
Theoretical fractional contribution to the overall lifetime. Comparing changes upon activation for a dye-based (blue) and FRET-based (red) biosensor. FRET biosensor: quenching of the donor fluorophore with 50% efficiency in the active (solid red) or inactive (dashed red) conformation. Dye-based biosensor: brightness ratios of free to bound form are 1.0 (solid blue) or 0.1 (dashed blue).

### *In Vivo* lifetime Imaging

NIH 3T3 mouse embryonic fibroblasts (MEFs) were microinjected with **mcFLIM 87** and imaged on a confocal FLIM microscope. FLIM allowed us to visualize both the distribution of the biosensor **(Figure 5a)** and the absolute concentration of active Cdc42 **(Figure 5b, Methods)**. We first determined phasor locations of the free and bound biosensors from the *in vitro* assay (***p_f_*** and ***p_b_*** respectively, solid circles). The increase in emission intensity produced by binding active Cdc42 resulted in a nonlinear relationship between the fraction of activated protein (hash marks, increments of 10%) and the fractional contribution of each species to the overall lifetime. Cells loaded with the wildtype biosensor **(Figure 5a-c, column 1-2)** had phasors that displayed a smear along the line connecting the two states, indicating a distribution of Cdc42 activity within the cells. The majority of pixels ranged between 0 and 10% of what would be observed for fully bound biosensor, as indicated by the first two hash marks. This was consistent with previous biochemical studies (Boulter et al., 2010; Jennings and Knaus, 2014). It is noteworthy that this relatively small change in binding produced a readily discernable change in lifetime represented by the phasor plot. In contrast, in the negative control cells where a mutation of the affinity reagent abolished the interaction with Cdc42 **(Figure 5a-c, column 3)**, the measured lifetimes were bunched near the free form of the biosensor, signifying negligible Cdc42 activation.

**Figure 5.**
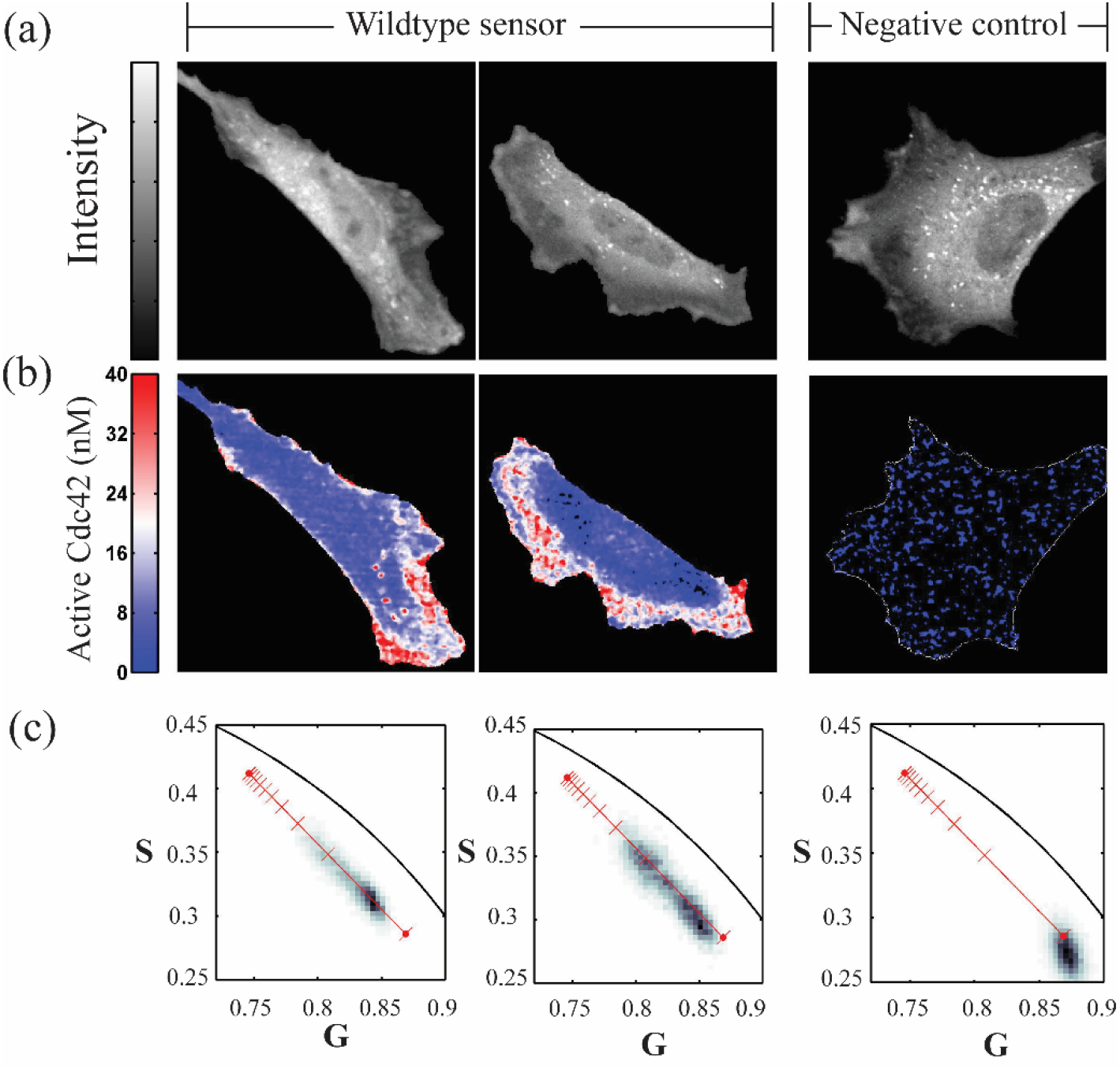
(**a**) Intensity, (**b**) activated cdc42 concentration and (**c**) phasor histogram for cells injected with **mcFLIM 87** or **mcFLIM 87 mutated to abrogate Cdc42 binding**. The phasor histograms are labeled with the locations of free and bound biosensor (solid red circles). The hash marks indicate 10% increments of protein activation.

Knowledge of active protein concentrations provides information valuable for the biochemical modeling of regulatory cycles. The spatial distribution of active Cdc42 concentrations across individual cells were calculated by analyzing phasors as described in the materials and methods **(Eq. 5)**. Shown in Fig. 5b are the pseudo-colored, mapped concentrations of active Cdc42. Endogenous Cdc42 activation was observed near protrusions, consistent with previous results (Itoh et al., 2002; Nalbant et al., 2004) **(Figure 5b)**. The relatively large dynamic range of lifetimes for small fractions of active proteins resulted from the increase in emission intensity upon protein binding.

## Conclusion

Dye-based FLIM biosensors, studied with phasor analysis, can provide a powerful tool for monitoring the conformations of proteins in vivo. Dye-based biosensors have not been widely used, likely because they formerly required isolation and labeling of proteins, followed by cumbersome procedures for loading them into cells. Recent advances enabling labeling within live cells could bring them into reach (e.g. unnatural amino acid incorporation (MacNevin et al., 2019) and self-labeling enzymes (Keppler et al., 2003; Los et al., 2008)). Here we evaluated a series of dyes, all of which showed promising brightness and solvent-induced FLIM changes, for use in FLIM biosensors. Although one dye was clearly superior for monitoring Cdc42, the others provide alternatives that can be used on affinity reagents that experience different ranges of solvent polarity or hydrogen bonding. The rigidity and length of the linker will be important in drawing the maximum response in each circumstance, positioning the dyes where their access to water or hydrogen-bonding residues is affected by target binding.

Past studies have focused on FRET-based biosensors, but biosensors based on merocyanines have enhanced capacity to report activity when low percentages of the target protein are activated. Furthermore, the biosensor design here responds to endogenous protein, so should be less perturbing than other approaches. Combining reduced perturbation with quantitation of low activated concentrations bodes well for multiplexing, the study of low abundance targets, and understanding phenomena where only small proportions of proteins are activated. We hope that the dyes and design described here can provide the advantages of dye-based FLIM biosensors to a broad range of biological studies.

## SUPPORTING MATERIAL

We gratefully acknowledge the NIH for their support of this work (NIGMS R35 GM122596 and NCI CA252826).

